# Redundant functions of the SLC5A transporters Rumpel Kumpel and Bumpel in ensheathing glial cells

**DOI:** 10.1101/2021.09.28.462185

**Authors:** Kerem Yildirim, Bente Winkler, Nicole Pogodalla, Steffi Mackensen, Marie Baldenius, Luis Garcia, Elke Naffin, Silke Rodrigues, Christian Klämbt

## Abstract

Neuronal processing is energy demanding, and relies on sugar metabolism as an energy source. To provide a constant metabolite supply neurons and glial cells express many glucose and lactate transporters of the solute carrier (SLC) 5A family. Here we dissect the partially redundant functions of three highly related glia specific Drosophila genes encoding SLC5A proteins, Rumpel, Bumpel and Kumpel. While knockdown of *rumpel* causes several behavioral phenotypes, they are less prominent in *rumpel* mutants. *bumpel* and *kumpel* mutants are viable and fertile, lacking discernible phenotypes. However, in *bumpel kumpel* double mutants and to an even greater extent in *rumpel bumpel kumpel* triple mutants oogenesis is disrupted at the onset of the vitollegenic phase. This indicates at least partially redundant functions between these genes. Rescue experiments exploring this effect indicate that oogenesis can be affected by CNS glial cells. Moreover, expression of heterologous mammalian SLC5A transporter proteins, with known transport properties, suggest that Bumpel and/or Kumpel transport glucose or lactate. Overall, our results imply a redundancy in SLC5A nutrient sensing functions in Drosophila glial cells, affecting ovarian development and behavior.

## Introduction

Cells require a constant energy supply to function. Metabolic activity is in particular high in the nervous system, where large amounts of ATP is needed to maintain synaptic transmission and cope with the resulting changes in membrane potential. This is reflected by energy consumption as the mammalian brain accounts for 20% of the total resting oxygen consumption although comprising only 2% of the body’s weight (Karbowski, 2007; Mink et al., 1981; Nortley and Attwell, 2017). This energy demand is even greater in young brains and is similarly found in the invertebrate nervous system (Harris et al., 2012; Laughlin et al., 1998; Mink et al., 1981; Tsacopoulos et al., 1988).

In the vertebrate nervous system, glucose is the predominant metabolite supplying the brain with energy. Glucose circulates in the blood stream and is delivered to the different organs. The brain is metabolically separated from circulation by the blood-brain barrier which is comprised of endothelial cells that form occluding tight junctions (Abbott et al., 2006; Tam and Watts, 2010; Tietz and Engelhardt, 2015; Zlokovic, 2008). Endothelial cells take up glucose from the blood stream via the Glut1 transporter. From the endothelial cells, glucose is then shuttled to astrocytes and neurons by different glucose transporters. While endothelial cells and astrocytes express differentially glycosylated forms of the glucose transporter Glut1, neurons predominantly express Glut3 (Barros et al., 2007; Vannucci et al., 1997). To match fluctuating neuronal energy demands, glial cells are able to sense synaptic activity. The Astrocyte Neuron Lactate Shuttle (ANLS) hypothesis, initially established for the mammalian brain provides an elegant model explaining how the flux of small C3 metabolites is regulated in the brain (Magistretti and Allaman, 2018; Pellerin and Magistretti, 1994; Pellerin et al., 2007).

In contrast to vertebrates, invertebrates do not have a vascular system. Instead, the hemolymph, the blood equivalent tissue of invertebrates, is found in all body cavities and immerses the entire nervous system. The predominant sugar in hemolymph is trehalose, a non-reducing disaccharide composed of two glucose molecules linked in an α,α-1,1-glycosidic manner, acting as the prime energy source (Wyatt and Kalf, 1957). As in vertebrates, the nervous system is metabolically separated from the remaining body by the blood-brain barrier (Carlson et al., 2000; Limmer et al., 2014; Mayer et al., 2009). In Drosophila the blood-brain barrier is established by perineurial and subperineurial glial cells (Stork et al., 2008). Perineurial cells express trehalose transporters and participate in maintaining the energy homeostasis of the brain (Volkenhoff et al., 2015). The subperineurial glial cells block paracellular diffusion by interdigitating cell-cell processes and the formation of septate junctions (Babatz et al., 2018; Bundgaard and Abbott, 2008; Schwabe et al., 2005; Stork et al., 2008). Trehalose is taken up from the circulation by the Tret1-1 transporter, which is expressed by perineurial glial cells. In addition, MFS3 and Pippin are involved in carbohydrate transport in the perineurial glia. Interestingly, MFS3 or Pippin null mutants are rescued via compensatory upregulation of Tret1-1, another BBB carbohydrate transporter, while RNAi-mediated knockdown of *Mfs3* and *pippin* is not compensated for (McMullen et al., 2020). Trehalose is subsequently metabolically processed through glycolysis. Lactate and alanine are then delivered to neurons by as yet poorly characterized transport mechanisms (Delgado et al., 2018; González Gutiérrez et al., 2019; Volkenhoff et al., 2015).

In general metabolite transport is mediated by members of the solute carrier protein (SLC) family, which allow either facilitated, or active transport into the cell. The SLC superfamily constitutes approximately 400 genes grouped into more than 50 families and many of its members are expressed in the brain (Bai et al., 2017; Morris et al., 2017). Two of these transporter families have been identified to be involved in glucose transport. The solute carrier proteins of the SLC2A family (Glut1-14) mediate facilitated glucose diffusion across the plasma membrane, whereas members of the SLC5A family (SGLT1-5) transport glucose in a sodium gradient-dependent manner (Mueckler and Thorens, 2013; Wright, 2013; Wright et al., 2011).

In Drosophila, it is long known from deoxyglucose labeling experiments that glucose can be taken up by neurons in the brain (Buchner et al., 1979). Moreover, recent experiments using a FRET-based glucose sensor expressed in neurons of the Drosophila CNS demonstrated that neurons are able to take up glucose in the same manner as glial cells (Volkenhoff et al., 2018). The fly orthologue of the mammalian Glut1 transporter is expressed exclusively in neurons and is not expressed in the blood-brain barrier (Volkenhoff et al., 2018).

The cellular route and transporters involved in delivery of trehalose, as well as glycolytic derived products to neurons remains elusive. The Drosophila nervous system comprises a relatively small set of well-defined glial cells, which establish a glial network that connects glial cells of the blood-brain barrier with the synaptic neuropil (Freeman, 2015; Yildirim et al., 2018). The cortex glial cells engulf neuronal cell bodies (Coutinho-Budd et al., 2017; Spéder and Brand, 2018). Axons and dendrites are located in the neuropil, which is infiltrated by numerous fine cell processes of the astrocyte-like glial cells (MacNamee et al., 2016; Peco et al., 2016; Stork et al., 2014). These cells modulate synaptic activity by participating in neurotransmitter homeostasis and the secretion of additional modulatory factors (Liu et al., 2014; Ma et al., 2016; Sengupta et al., 2019). The cell bodies of the astrocyte-like glia cells are found at the boundary of the neuropil, next to the ensheathing glial cell bodies (Peco et al., 2016). Ensheathing glia encase the entire neuropil and also participate in the modulation of locomotor activity, as well as in the regulation of sleep (Otto et al., 2018; Stahl et al., 2018).

Here we used Drosophila to uncover further glial transporters required for maintaining neuronal function. In a cell type specific RNAi based screen (Schmidt et al., 2012) we identified several genes that when silenced in glial cells cause abnormal adult behavior. One such gene is *CG9657*, which encodes a SLC5A family member. Due to the locomotion defect observed following glial specific gene silencing, the gene was named *rumpel* according to the Sesame street character who barely moved. In contrast to knockdown animals, *rumpel* mutants show only very subtle behavioral phenotypes suggesting compensatory genetic mechanisms. We therefore analyzed the SLC5A gene family for the closest homologs and identified *kumpel* and *bumpel* (for *kin of rumpel* [*kumpel*] and *brother of rumpel* [*bumpel*]). These two genes are expressed partially overlapping with *rumpel* and single mutants show no severe phenotypes. A triple mutant affecting *rumpel*, *kumpel* and *bumpel*, however, showed behavioral phenotypes and female sterility demonstrating redundant gene functions.

## Results

### Identification of *rumpel*

*CG9657* was identified in an RNAi-based screen for adult locomotor deficits using a construct without any predicted off-target (Dietzl et al., 2007; Schmidt et al., 2012). In adult flies, pan-glial suppression of *CG9657* results in a bang sensitive paralysis phenotype. Upon 10 sec of vortexing, wild type flies immediately recover and show normal locomotor activity. In contrast, *CG9657* knockdown flies are paralyzed for approximately 10 minutes before regaining consciousness and flying away. In addition to mechanical stimuli, high temperature (37°C) triggers paralysis of these flies in the same manner. In a next step, we analyzed the effects of glial *CG9657* knockdown on larval locomotion using FIM (Risse et al., 2017b; Risse et al., 2013). When placed on the warmest side of the table, with a temperature gradient ranging from 32°C to 16°C, control larvae quickly move towards cooler temperature (**Fig. 1A,B**). In contrast, upon glial knockdown of *CG9657*, animals showed very little movement at elevated temperatures, however were able to respond to the temperature gradient when placed at intermediate temperatures (**Fig. 1C,D**). Based on the slow-moving character of the Sesame street, we named the gene as *rumpel*.

**Figure 1.**
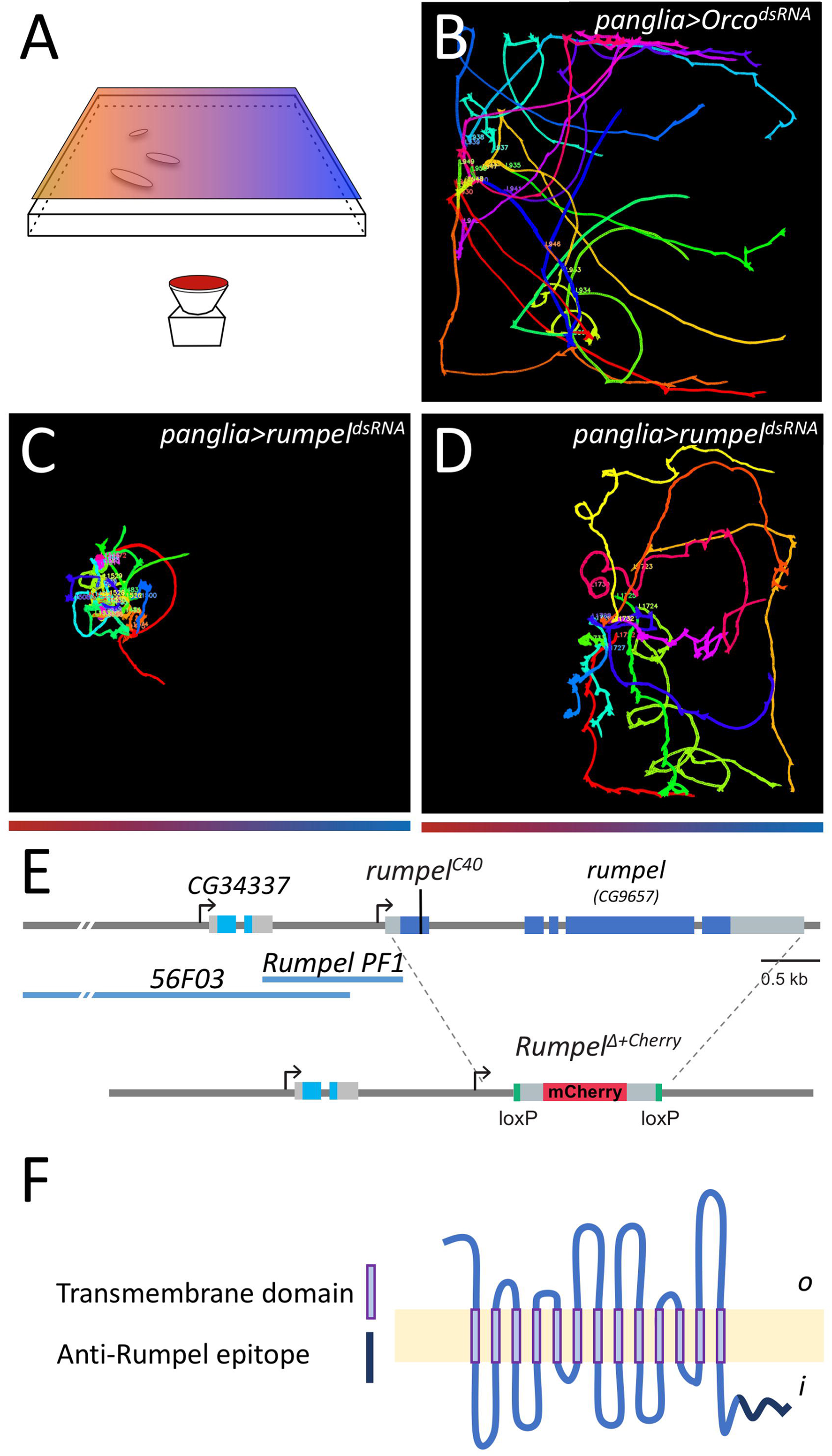
RNAi based suppression of *rumpel* results in temperature dependent paralysis. **(A)** To test whether temperature affects larval locomotion we established a temperature gradient from 32°C to 16°C (from left to right) on a 25 cm x 25 cm large tracking arena. Larval locomotion is analyzed using FIM. **(B)** 10 third-instar larvae of the wandering stage with panglial expression of control dsRNA (*Orco^100825^*) were placed at 32°C on the tracking arena and imaged for 10 minutes. Control larvae crawl towards cooler temperatures. **(C)** 10 third-instar larvae of the wandering stage with pan-glially reduced *rumpel* expression [repo-Gal4, *UAS-rumpel^dsRNAv43922^*] were placed at 32°C. Note, larvae do not move towards cooler temperatures. **(D)** 10 third-instar larvae of the wandering stage with panglially reduced *rumpel* expression placed at the 28°C position are able to detect the temperature gradient and move towards cooler temperatures. **(E)** Schematic representation of the *rumpel* (*CG9657*) locus on the X-chromosome. Exons are shown in boxes, *rumpel* coding exons are in dark blue, *56F03* and Rumpel *PF1* denote enhancer elements that direct expression in ensheathing glia. The position of the CRISPR-induced premature stop codon in amorphic allele (*rumpel^C40^*) and the *rumpel* locus replacement with *attP-loxP-Cherry-loxP* in (*rumpel^Δ+Cherry^*) is indicated. **(F)** The Rumpel protein is predicted to have 13 membrane (light yellow) spanning domains. The peptide sequence used to immunize rabbits is highlighted in dark blue. o: outside, i: inside.

To determine in which glial cells *rumpel* is required, we performed knockdown experiments using glial subtype specific Gal4 drivers. Silencing in perineurial, subperineurial or cortex glial cells (*apontic-Gal4, moody-Gal4* or *55B12-Gal4*, respectively) did not affect locomotor abilities of third instar larvae. Suppression of *rumpel* in neuropil-associated glial cells, astrocyte-like cells and ensheathing glial cells (*nrv2-Gal4, alrm-Gal4* or *83E12-Gal4*, respectively), however, caused a temperature induced paralysis phenotype. Thus, *rumpel* appears to be required in the neuropil associated glial cells.

### *rumpel* is expressed by ensheathing and astrocyte-like glia

The *rumpel* gene is situated on the X-chromosome and encodes a predicted sugar transporter protein of the sodium solute symporter 5A (SLC5A) family with 13 transmembrane domains (**supplementary Fig. 1; Fig. 1E,F**). To further identify the cells expressing *rumpel* we dissected the *rumpel* promotor region. A 1.1 kb long enhancer fragment designated as *rumpel^PF1^* (**Fig. 1E**) directs specific expression in the nervous system only in cells that are Repo positive (**Fig. 2A,B**). Based on their location around the neuropil, the *rumpel* expressing cells may correspond to ensheathing glial cells and/or astrocyte-like glial cells. To further test which cell type activates the *rumpel* enhancer we crossed the *rumpel^PF1^-stGFP* construct into a genetic background directing the expression of a red nuclear marker in the ensheathing glial cells [*rumpel^PF1^-stGFP*, *nrv2-Gal4*; *UAS-stRed*] (**Fig. 2C**). Most *rumpel* expressing neuropil-associated cells also show *nrv2-Gal4* activity, suggesting that *rumpel* positive cells are expressed in ensheathing glial cells. This notion is corroborated by split-Gal4 experiments where we co-expressed the Gal4-DNA-binding domain in the *rumpel* pattern and the Gal4 activation domain in the *nrv2* pattern [*rumpel^PF1^-Gal4^DBD^*, *nrv2^PF4^-Gal4^AD^*] (**Fig. 2D**). To test whether Rumpel is also expressed by astrocyte-like glial cells, we analyzed animals expressing GFP under the control of the *rumpel* enhancer and dsRed under the control of the *alrm* enhancer, which is active in astrocytes. In the larval central nervous system of such animals we noted frequent coexpression (**Fig. 2E**), suggesting that Rumpel is also expressed by astrocyte-like glial cells. Similarly, when we stained *rumpel^PF1^-stGFP* larval brains with anti-Nazgul antibodies we noted a partial overlap (**Fig. 2F**). The *rumpel^PF1^* fragment overlaps with the enhancer fragment *56F03* generated by the FlyLight project (Jenett et al., 2012; Li et al., 2014), which is reported to direct expression in ensheathing glia (Li et al., 2014; Otto et al., 2018; Peco et al., 2016) (**Fig. 1E**). This indicates that the critical enhancer elements are located in the 700 bp overlap of the two enhancer fragments.

**Figure 2.**
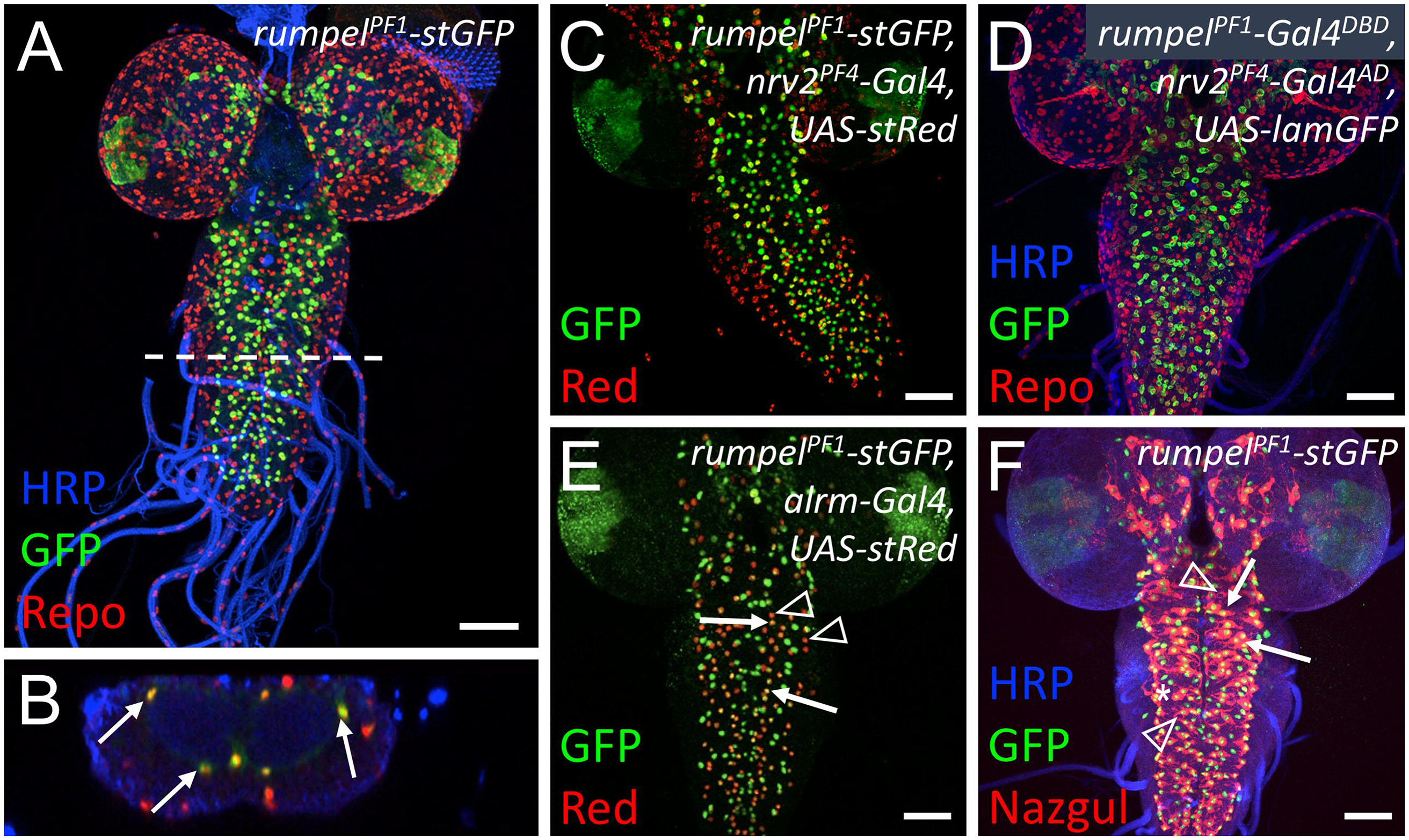
*rumpel*-PF1 induces an expression in the neuropil-associated glial cells. **(A-C,E-F)** Specimens are stained for promoter fragment induced expression of StingerGFP (stGFP, green). **(C,E)** RedStinger (stRed, red). **(D)** LaminGFP (lamGFP, green). **(A,D)** Glial nuclei are stained for Repo protein localization (red). **(F)** Astrocyte-like glial cells are stained for Nazgul protein localization (red). Neuronal membranes are shown in blue (HRP staining). **(A)** *rumpel* promoter fragment PF1 (*rumpel^PF1^*) induces stGFP expression in Repo positive cells in the third instar larval brain. White dashed line indicates the position of the orthogonal section shown in **(B)**. **(B)** Glial cells in the position of ensheathing glia are indicated by arrows. No expression is observed in surface associated glial cells. **(C)** *rumpel^PF1^* induced stGFP expression overlaps with the nrv2 induced RedStinger expression. **(D)** Split Gal4 directed expression of LamGFP is found in ensheathing glial cells [*rumpel^PF1^-Gal4DBD*, *nrv2PF4-Gal4^AD^*, *UAS-lamGFP*]. **(E)** *rumpel^PF1^* induced stGFP expression is found in some astrocyte-like glial cells labelled by *alrm* induced stRed expression (compare arrows with arrowheads). **(F)** *rumpel^PF1^* induced stGFP expression in Nazgul positive astrocyte-like glial cells (arrows). The asterisk denotes ensheathing glial nuclei, the arrowhead denotes astrocytes not activating the *rumpel^PF1^* enhancer. Scale bars are 50 μm.

### Rumpel protein expression in the nervous system

To determine the localization of the Rumpel protein, we generated an anti-peptide antiserum directed against the C-terminal most amino acids (**Fig. 1F**). The specificity of the antiserum is demonstrated following pan-glial silencing of *rumpel* expression using RNAi (**Fig. 3A,B**). In third instar larvae, no expression is discernible outside the central nervous system, which matches RNAseq expression data (Brown et al., 2014; Graveley et al., 2011). Within the nervous system, Rumpel localizes to cell membranes of neuropil associated cells in the developing brain lobes as well as in the ventral nerve cord (**Fig. 3A-F, arrows**). In addition, some Rumpel protein can be found within the neuropil (**Fig. 3E,F, arrowheads**). Very low levels of Rumpel protein are detected along the peripheral abdominal nerves that connect the CNS with the periphery. In adults, Rumpel expression is also found prominently in the ensheathing glial cells (**Fig. 3G,H**).

**Figure 3.**
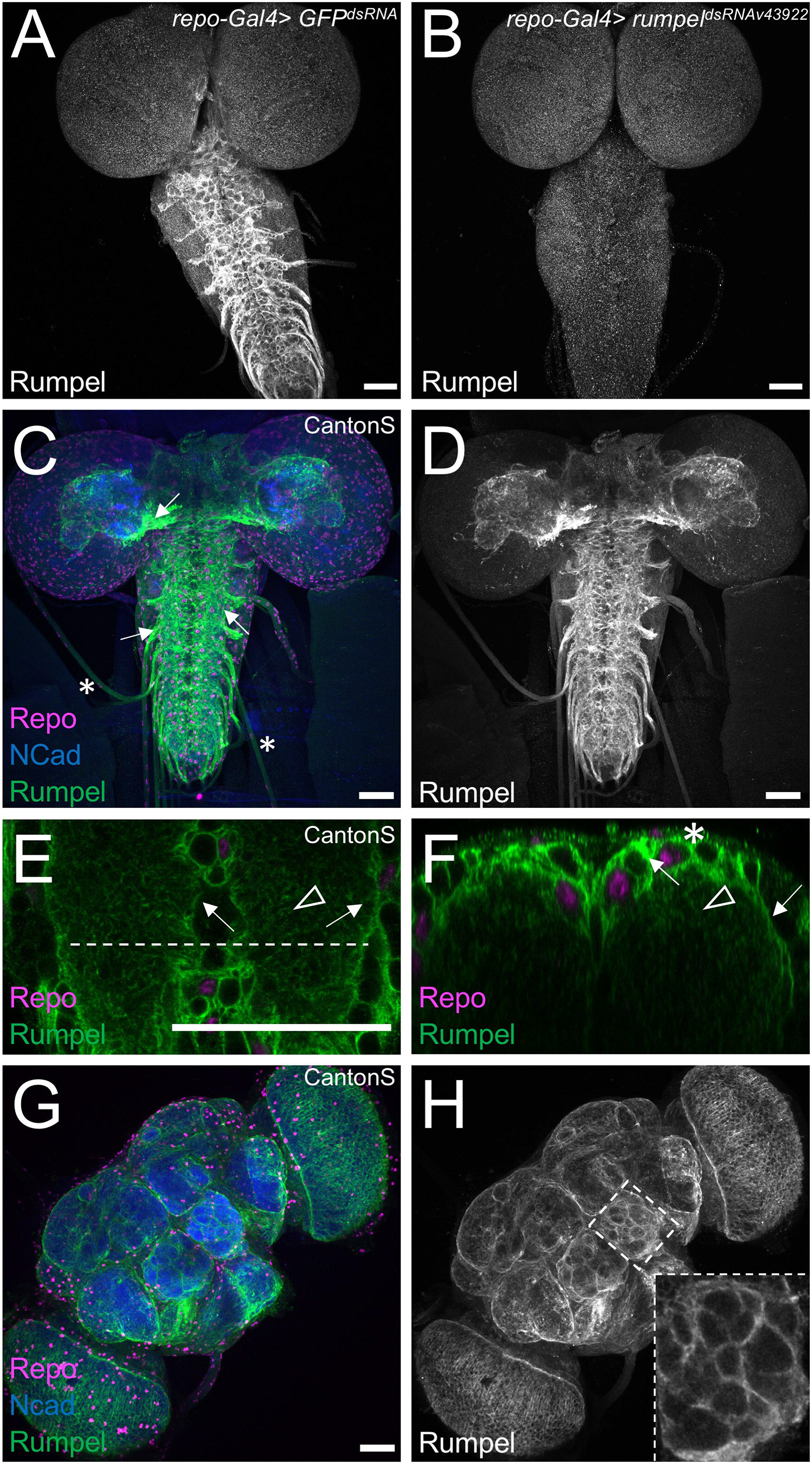
Rumpel protein is expressed in the neuropil-associated glial cells. All specimens are stained for Repo localization to define glial nuclei (magenta), for N-Cadherin localization to visualize axonal and dendritic cell membranes (blue) and for Rumpel protein localization (green/grey). **(A-F)** Third instar larval brains and **(G,H)** adult brain. **(A)** In control animals [*repo-Gal4*, *UAS-GFP^dsRNA^*] Rumpel protein localizes around the neuropil. **(B)** Upon expression of *rumpel^dsRNA^* in the all glial cells [repo-Gal4, *UAS-rumpel^dsRNAv43922^*] no Rumpel protein can be detected, demonstrating the specificity of the anti-Rumpel antibody. **(C,D)** Rumpel localization is observed surrounding the neuropil (arrows) in a position of the ensheathing glial cells. Very little Rumpel protein is found along larval nerves (asterisks). **(E)** Image of a single confocal plane through a third instar larval ventral nerve cord. Rumpel localizes to ensheathing glial cell membrane (arrow) and to cell processes of astrocyte-like glial cells (arrowhead). The dashed line indicates the position of the orthogonal section shown in **(F)**. **(F)** Rumpel localizes to ensheathing glial cells (arrows) and astrocytic processes in the neuropil (arrowhead). Note, the pronounced cortex-glial cell like ramifications of the ensheathing glia dorsally to the neuropil (asterisk) **(G,H)** Rumpel localizes around the neuropil in adult brains at a position of the ensheathing glia (inset: antennal lobe). Scale bars are 50 μm.

To further characterize the Rumpel expressing glial cells, we performed glial cell type specific silencing experiments. Following suppression of *rumpel* using *nrv2-Gal4*, which strongly suppresses in ensheathing glial cells and less so in cortex and astrocyte-like glial cells, we noted a complete lack of Rumpel protein localization (**Supplementary Fig. 2A,D**). Following suppression in ensheathing glial cell using *83E12-Gal4*, weak Rumpel expression can be detected in the cortex and neuropil, reflecting processes of the astrocyte-like glial cells (**Supplementary Fig. 2B,E**). Following suppression of *rumpel* expression, mostly in astrocyte-like glial cells, using *alrm-Gal4*, Rumpel protein can still be detected in the ensheathing glia (**Supplementary Fig. 2C,F**).

Similar to that of the larval nervous system, Rumpel protein is also primarily detected in ensheathing glial cells of the adult CNS (**Fig. 3G,H**). This is also corroborated by recent single cell RNA sequencing data (Davie et al., 2018) (**supplementary Fig. 3A-I**). Expression of the glial cell marker Repo defines all glial cells in the adult fly brain (Halter et al., 1995). The different glial subtypes are characterized by expression of specific genes [perineurial glial cells: *tret1-1* (Volkenhoff et al., 2015), subperineurial glial cells: *gliotactin* (Auld et al., 1995; Babatz et al., 2018), cortex glia: *zydeco* (Melom and Littleton, 2013), astrocyte-like glial cells: *GAT* and *nazgul* (Ryglewski et al., 2017; Stork et al., 2014) and ensheathing glial cells: EAAT2 (Peco et al., 2016)]. The different glial subtypes cluster in distinct groups of cells (Davie et al., 2018) (**supplementary Fig. 3A-I**). In adult glial cells, Rumpel is expressed most strongly in the ensheathing glia cluster but in addition some cortex glia and astrocyte-like glial cells express *rumpel* (**Fig. 3G,H; supplementary Fig. 3A-I**).

In conclusion, throughout development of the central nervous system of Drosophila, the SLC5A member Rumpel is expressed specifically in glial cells and is most prominently found in ensheathing glial cells with some expression in cortex and astrocyte-like glial cells.

### Generation of *rumpel* mutants

The analysis of Rumpel protein localization demonstrates a specific expression in neuropil associated cells. Moreover, the RNAi data suggests a role of *rumpel* in neuropil-associated glial cells, to control neuronal activity during noxious stimuli such as heat and mechanical stress. To further validate these phenotypic findings, we generated a loss-of-function allele using the CRISPR/Cas9 methodology, targeting either the first or the fourth exon. The allele *rumpel^C40^* carries a 7 bp deletion leading to a frameshift and subsequent stop of translation after 20 additional amino acids (**Fig. 1E**). In addition, we replaced the entire *rumpel* gene with an *mCherry* coding sequence using homologous recombination to generate the *rumpel^Δ+cherry^* mutant (see Materials and Methods for a description of all mutants) (**Fig. 1E**). Both *rumpel* null alleles generated lack detectable expression of the Rumpel protein. Moreover, both *rumpel* null alleles are homozygous viable and show no discernible morphological phenotypes. Likewise, a floxed allele, *rumpel^del^*, shows no detectable phenotypic abnormalities.

### Behavioral analysis of *rumpel* null mutants

RNA interference based knockdown of *rumpel* caused a prominent paralysis of the adult flies upon mechanical stress or elevated temperature. To our surprise, *rumpel* null mutant flies fail to show any of these responses and behaved as wild type flies. To better quantify behavioral phenotypes we turned to larval locomotion. RNAi based knockdown of *rumpel* causes a temperature dependent locomotion phenotype (**Fig. 1**). This was not observed for mutant *rumpel^C40^* larvae. However, when comparing unconstrained locomotion at 25°C and at 32°C we noted slight differences. A heat map representation of control and *rumpel* mutant larvae shows that at 25°C both control as well as *rumpel* mutants spread evenly across the tracking arena (**Fig. 4A,B**). In contrast at 32°C, *rumpel^C40^* null mutants do not explore the tracking plate as intensively as control larvae (**Fig. 4C,D**). This reduced exploratory locomotion phenotype is reflected in the mean distance to origin of the mutant animals (**Fig. 4E**, n=150 larvae, 3 minutes tracking, p = 0.023). Interestingly, when we tested *rumpel^Δ+cherry^*, that was backcrossed 10 times against a *w^1118^* background, we noted no significant change in distance to origin at elevated temperatures (**supplementary Fig.4**). Ensheathing glial cells have been associated with sleep phenotypes (Davla et al., 2020; Stahl et al., 2018). Thus, we tested whether *rumpel* null mutants show an abnormal sleep behavior. We noted a significant increase in day sleep compared to *w^1118^* flies, similar to what was observed for taurin transporter (EAAT2) (Davla et al., 2020; Stahl et al., 2018) (**Fig. 4F,G**).

**Figure 4.**
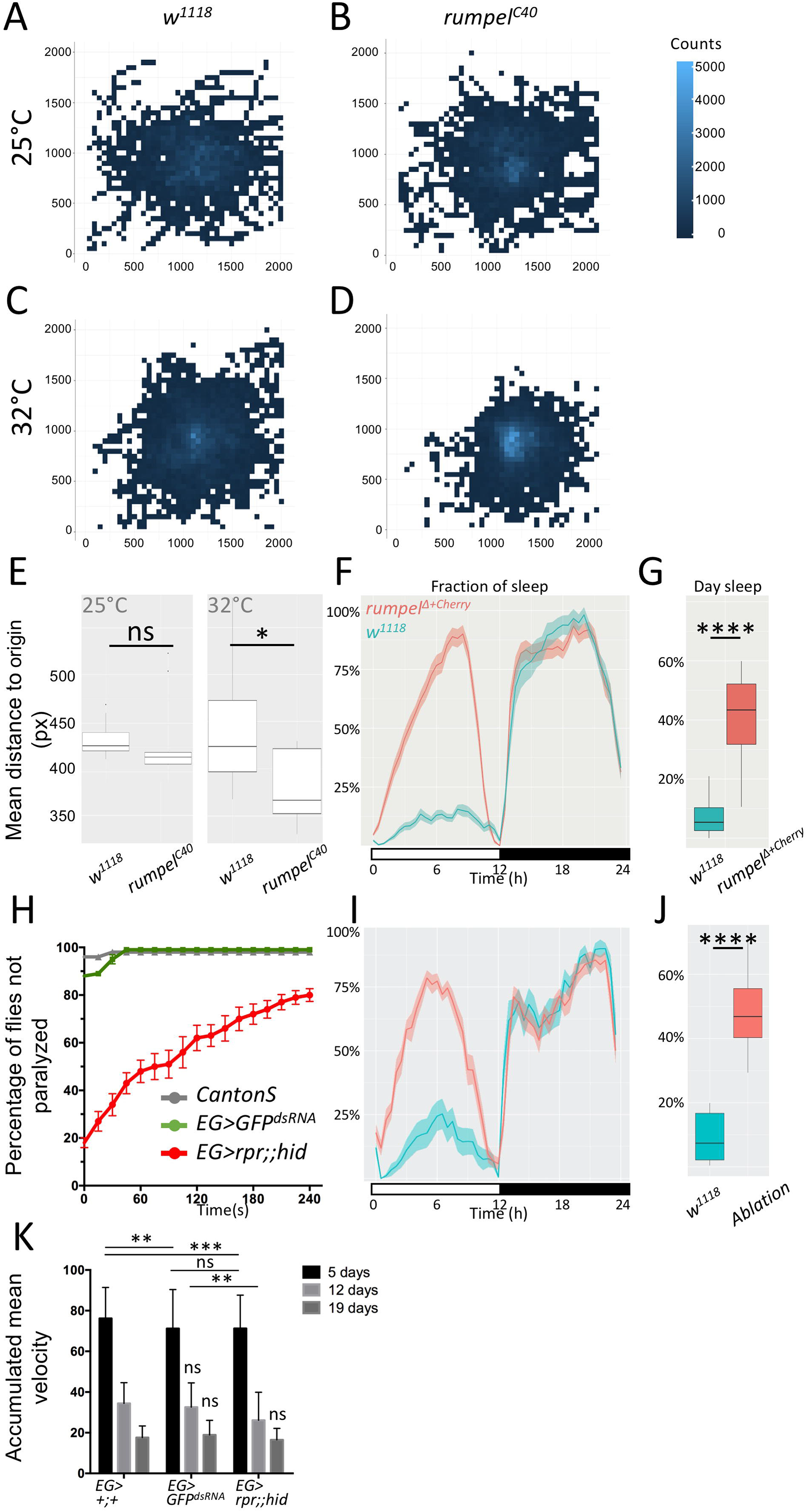
*rumpel^C40^* has a temperature dependent larval locomotion phenotype. **(A-E)** 150 third instar larvae of the respective genotypes were recorded in groups of 15 animals for three minutes at 25°C or 32°C, as indicated. Larvae were always placed at the middle of the tracking plate. **(A-D)** For heatmap analyses, the 2048×2048 px image of the agar plate is divided in 50×50 px squares. The number of larval appearances per square is determined and indicated in blue shading using R. Darker blue colors indicate less frequent appearance, while lighter blue ones more. **(A)** Heatmap analysis of control *w^1118^* larvae at 25°C. Wild type larvae crawl in every direction and spread evenly on the agar plate at 25°C (indicated by fewer lighter blue squares). **(B)** Heatmap analysis of *rumpel^C40^* larvae. *rumpel^C40^* larvae shows wild type like distribution on the agar plate at 25°C (indicated by similar number of lighter blue squares). **(C)** Wild type larvae spread evenly on the agar plate at 32°C. **(D)** At 32°C *rumpel^C40^* larvae spread less on the agar plate (indicated by more light blue squares in the middle). **(E)** Quantification of the mean distance to origin of wild type vs *rumpel^C40^* larvae at 25°C and 32°C. At 25°C no significant difference is indicated (Mann-Whitney U Test p > 0.05, n=150). Mean distance to origins of wild type and *rumpel^C40^* are 439.4 and 381.4 px, respectively at 32°C. Wild type larvae spread significantly more on the agar plate at 32°C compared to *rumpel^C40^* larvae (Mann-Whitney U Test p = 0.023, n=150). **(F)** To monitor the effects of rumpel on sleep behavior *rumpel^Δ+cherry^* flies were backcrossed 10 times to *white^1118^*. The activity of 40 flies was tracked over 7 days in the ethoscope (Geissmann et al., 2017). **(G)** *rumpel^Δ+cherry^* flies sleep significantly more during the day (p = 2e-16, Wilcoxon rank sum), whereas night sleep is not affected. **(H)** Heat shock assay of 5 days old male and mated female flies of wildtype [*Canton S*], [*GMR83E12-Gal4^AD^*; *repo-Gal4^DBD^*, *UAS-GFP^dsRNA^*], [*UAS-rpr*; *GMR83E12-Gal4^AD^*; *repo-Gal4^DBD^*, *UAS-hid*] (each genotype n=100). Flies are heat shocked in a water bath for 2 min at 40 °C and were immediately recorded at room temperature. Not moving flies lying on their back are considered as paralyzed. Recording was stopped after 240 sec. Error bars indicate standard deviation. **(I)** Average sleep time over seven days summarized for 24 h (shown in %). Flies lacking ensheathing glia show an increased day time sleep compared to the control. **(J)** Summary of the fraction of time sleeping over seven days (shown in %). Loss of ensheathing glia leads to an increased sleeping time during the day (p- = 9.991e-07, Wilcoxon rank sum). **(K)** The rapid iterative negative geotaxis (Ring) assay (Gargano et al., 2005) shows the climbing ability of females with the genotypes: [*GMR83E12-Gal4^AD^*; *repo-Gal4^DBD^*], or [*GMR83E12-Gal4^AD^*; *repo-Gal4^DBD^*, *UAS-GFP^dsRNA^*], or [*UAS-rpr*; *GMR83E12-Gal4^AD^*; *repo-Gal4^DBD^*, *UAS-hid*]. The age of tested flies is indicated. Flies are heat shocked in a water bath for 2 min at 40 °C and immediately recorded at room temperature for 240 sec. Non-moving flies lying on their back are considered to be paralyzed. Both control and ensheathing glia ablated flies show a similar age-related decline of locomotor abilities. p-values are: 5 day old flies: p _pEG>+ / EG>GFPdsRNA_ = 0.0014, p _pEG>+ / EG>rpr,hid_ = 0.0009, p _EG>GFPdsRNA / EG>rpr,hid_ > 0.9999, 12 days old flies: p _EG>GFPdsRNA / EG>hid_ = 0.0043. All other p-values are >0.05 = non significant (ns). Error bars indicate standard deviation. Quantification was done using a two way Anova multiple comparison.

*rumpel* null mutants do not become paralyzed at elevated temperatures unlike RNAi knockdown larvae. A point mutation leading to a truncation of the open reading frame results in a reduced locomotor activity at elevated temperatures, whereas a *rumpel* deletion mutant shows no abnormal locomotor phenotype.

Rumpel is specifically expressed by ensheathing glia, therefore we tested the behavioral phenotypes of animals lacking ensheathing glia. Following expression of the pro-apoptotic genes *reaper* and *hid* specifically in ensheathing glia, these cells die and adult flies survive with a median longevity of 32 days instead of 53 days (Pogodalla et al., 2021). To test whether flies lacking ensheathing glial cells are susceptible to temperature shock we treated 5 days old male, and mated female flies for 2 minutes at 40°C in a water bath. Wild type Canton S flies as well as flies expressing *GFP^dsRNA^* recovered very quickly, and after 1 minute, all flies were moving again (**Fig. 4H**). Flies lacking ensheathing glia remained paralyzed for several minutes, resembling the *rumpel* RNAi knockdown phenotype. Even after 4 minutes, 20% of the ensheathing glia ablated flies remained paralyzed (**Fig. 4H**). When we compared sleeping behavior of *rumpel* mutants and ensheathing glial ablated flies we noted a similarly significant increase of daytime sleep in both genotypes (**Fig. 4F,G,I,J**). To test whether adult locomotor ability is generally affected, we also performed a rapid iterative negative geotaxis assay (RING assay) (Gargano et al., 2005). We separately analyzed 5, 12, and 19 days old females. 5 days old control flies harboring only the split Gal4 construct performed slightly better when compared to flies that express an GFP^dsRNA^ (**Fig. 4K**). Comparing flies expressing GFPdsRNA in ensheathing glia with those that lack ensheathing glia behave similar. Likewise, we noted similar locomotor abilities in aged flies. indicating that ensheathing glial cells do not equally affect all adult locomotor behavior (**Fig. 4K**).

Taking the differential phenotypes of the different *rumpel* mutants, RNAi experiments and the results of the ablation experiments into account, we propose this difference could be due to redundancy, therefore we searched for genes with a similar sequence homology to *rumpel*.

### Identification of Kumpel and Bumpel

*rumpel* encodes a predicted monocarboxylate transporter of the SLC5A family. The Drosophila genome encodes 15 members of this family (Featherstone, 2011) (**supplementary Fig. 3O**). Of these, CG6723 encodes a protein that shares 52% amino acid identity with the Rumpel protein (**Fig. 5A, supplementary Fig. 5**). We thus named CG6723 as *brother of rumpel* (*bumpel*). The gene *CG42235* shows a more complex organization and encodes five different SLC5A like proteins (**Fig. 5B**). These different isoforms share 43% to 48% amino acid identity with Rumpel and Bumpel (**supplementary Figure 5**). We therefore named this gene *kin of rumpel* (*kumpel*).

**Figure 5.**
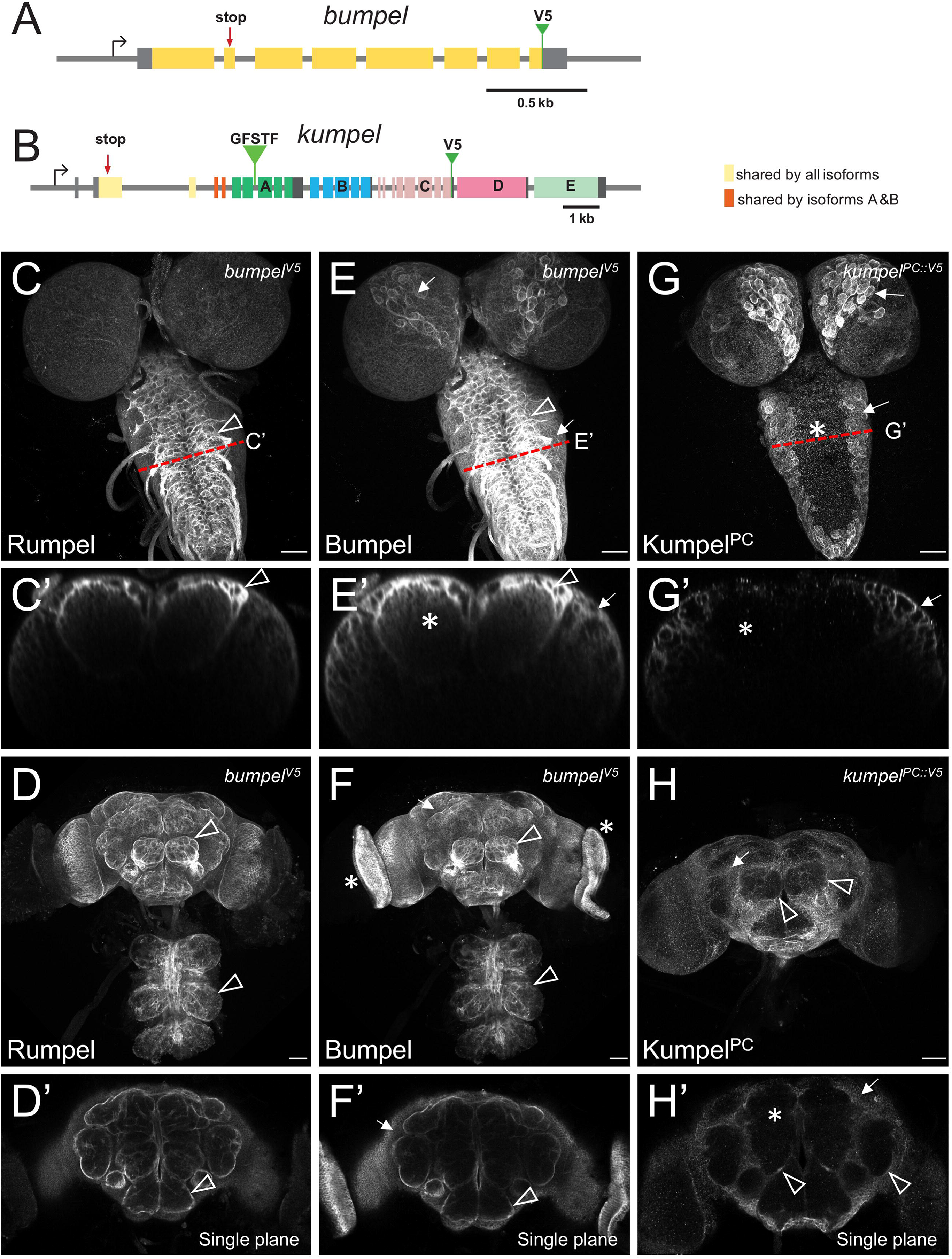
Bumpel and Kumpel are both expressed in CNS glial cells. **(A,B)** Schematic representation of the genomic loci of *bumpel* (*CG6723*) and *kumpel* (*CG42235*). Transcription is from left to right, coding exons are colored, five different isoforms are generated from the *kumpel* gene. The position of the stop codon mutations and the endogenously integrated V5 tags are indicated. GFSTF indicates the position of a MiMIC insertion. **(C-H)** Confocal analysis of third instar larval brains and adult brains stained for Rumpel, Bumpel^V5^ and Kumpel^PC∷V5^ as indicated. Red dotted lines indicate the position of orthogonal planes shown in (C’,E’,G’). **(C,C’)** Rumpel localizes predominantly in the ensheathing glial cells (arrowhead). **(D)** Maximum projections and **(D’)** single focal plane showing Rumpel localization in the adult brain. Rumpel is enriched in ensheathing glia (arrowheads). **(E,E’)** Bumpel^V5^ localizes to ensheathing glia (arrowhead) and cortex glial cells (arrows). Additional expression is noted in the neuropil (asterisk). **(F,F’)** In the adult nervous system, Bumpel localizes as detected for Rumpel. In addition, Bumpel^V5^ is found in the developing eyes (asterisk). **(G,G’)** Kumpel^PC∷V5^ localizes predominantly to cortex glial cells (arrows). No Kumpel^PC∷V5^ can be detected in the neuropil (asterisk). **(H,H’)** Kumpel^PC∷V5^ localizes to the cortex glial cells in the adult brain (arrows). Only weak expression in adult ensheathing glia is noted (arrowhead, H). No Kumpel^PC∷V5^ can be detected in the neuropil (asterisk).

### Rumpel and Bumpel share similar expression patterns

We next wanted to address whether *rumpel*, *bumpel* and *kumpel* are expressed by overlapping sets of glial cells. In situ hybridizations performed by the Berkeley genome project (Tomancak et al., 2002) showed also almost identical expression patterns of *bumpel* and *rumpel* during embryonic development (**supplementary Fig. 5**). Also for adult brains, single cell RNA sequencing data (Davie et al., 2018) suggest that *bumpel* is expressed in a very similar pattern as *rumpel* and is strongly expressed in ensheathing and weakly in astrocyte-like glial cells and cortex glial cells. In contrast, in adult brains *kumpel* expression is restricted to the ensheathing glia and not found in astrocytes (**supplementary Fig. 3I-K**).

To determine the protein localization of Bumpel, we inserted DNA sequences encoding a V5 tag at the 3’ end of the Bumpel coding region using a CRISPR-aided homologous recombination approach (see Materials and Methods for details, **Fig. 5A**). Homozygous *bumpel^V5^* flies eclosed in the expected Mendelian numbers and no abnormal phenotypes were detected. Endogenously tagged Bumpel protein is localized in a very similar pattern in the larval and adult brain, as observed for Rumpel (**Fig. 5C-F**). However, localization of Bumpel in cortex glial cells appeared slightly more pronounced (**Fig. 5C,E**). We also generated a *bumpel* minigene. *bumpel* is closely flanked by the genes *CG45676* and *Ipo9*. We cloned the entire *bumpel* gene locus including all flanking DNA sequences and the untranslated regions of *CG45676* and *Ipo9* and inserted a V5 tag at the C-terminus. The construct was placed on the second chromosome using the landing site 44F (Bischof et al., 2007) (**supplementary Fig. 6A**). In third instar larval brains, this construct also directs expression of Bumpel^V5^ in ensheathing and cortex glia (**supplementary Fig. 6B,C**).

### Kumpel expression in glial cells

In contrast to *rumpel* and *bumpel*, *kumpel* has a complex genomic organization and differential splicing is expected to generate five distinct isoforms (**Kumpel^PA-PE^, Fig. 5B**). Only the first two exons, which encode the N-terminal 236 amino acids are shared by all isoforms. These two exons encode the signal sequence and the first 1.5 of 13 predicted transmembrane domains. Two isoforms (PA and PB) share another two exons which encode a further 1.5 transmembrane domains. All other protein parts are unique to the different isoforms and given the conserved exon-intron arrangement, appear to originate from an ancient gene duplication.

To determine the *kumpel* expression pattern we inserted a V5 tag into the endogenous gene locus at the 3’ end of the last Kumpel^PC^ encoding exon (**Fig. 5B**). This isoform mostly localizes in cortex glial cells in the larval, as well as in the adult nervous system (**Fig. 5G,H**). Kumpel^PC^ appears to be weakly expressed by the adult ensheathing glia (**Fig. 5H, arrowheads**). Single cell sequencing data (**supplementary Fig. 3K**) indicates strongest expression of *kumpel* in ensheathing glial cells. Thus, other Kumpel isoforms may possibly show a more defined localization in ensheathing glial cells. To address their expression, we used an available converted MiMIC insertion line (*MI05542-GFSTS.0*) which directs expression of Kumpel^PA^-GFP fusion protein. However, this protein trap also labels mostly cortex glial cells (**supplementary Fig. 6D,E**).

### Generation of *bumpel* and *kumpel* mutants

To further study possible genetic relationships between *rumpel*, *bumpel* and *kumpel* we generated CRISPR induced mutants. The *bumpel* gene was targeted in first exon resulting in a frameshift at position +40 bp of the open reading frame, causing an early stop codon (**Fig. 5A**). The mutant is therefore predicted to be a null allele. Homozygous *bumpel* mutant flies eclosed at the expected Mendelian ratio and showed no fertility or morphological abnormalities. Adult flies also do not show any heat or bang sensitivity. Likewise, locomotion of *bumpel* mutant larvae is indistinguishable from the control at 25°C as well as at 32°C.

To induce *kumpel* mutants, we targeted the first common exon present in all *kumpel* transcripts (**Fig. 5B**). The mutation at position +253 bp of the reading frame caused a frameshift and subsequent termination of translation before the first transmembrane. The mutation can therefore be considered as null mutation. Mutant *kumpel* flies are homozygous viable and fertile and show no discernible abnormal phenotypes. As noted for *bumpel* mutant flies they show no locomotor deficits.

### *rumpel, bumpel* and *kumpel* genetically interact

*rumpel, bumpel* and *kumpel* encode highly related proteins that show similar expression patterns. To further determine possible redundancy between the different gene functions we first generated double mutant combinations. *rumpel bumpel* or *rumpel kumpel* double mutants are viable and fertile. Larval locomotion of double mutants is as of the single mutants. *bumpel kumpel* double mutant flies are also viable. However, homozygous females show reduced egg laying and are sterile. In these double mutants, oogenesis initially proceeds normally until stage eight. However, during the subsequent vitellogenic phase oogenesis appears disrupted and no normal eggs are formed, and they cannot be fertilized (**Fig. 6A-D**). We next generated *rumpel bumpel kumpel* triple mutant flies. Triple mutant females are sterile and do not lay any eggs. In contrast to the *bumpel kumpel* double mutant, oogenesis appears completely blocked after stage eight (**Fig. 6E,F**).

**Figure 6.**
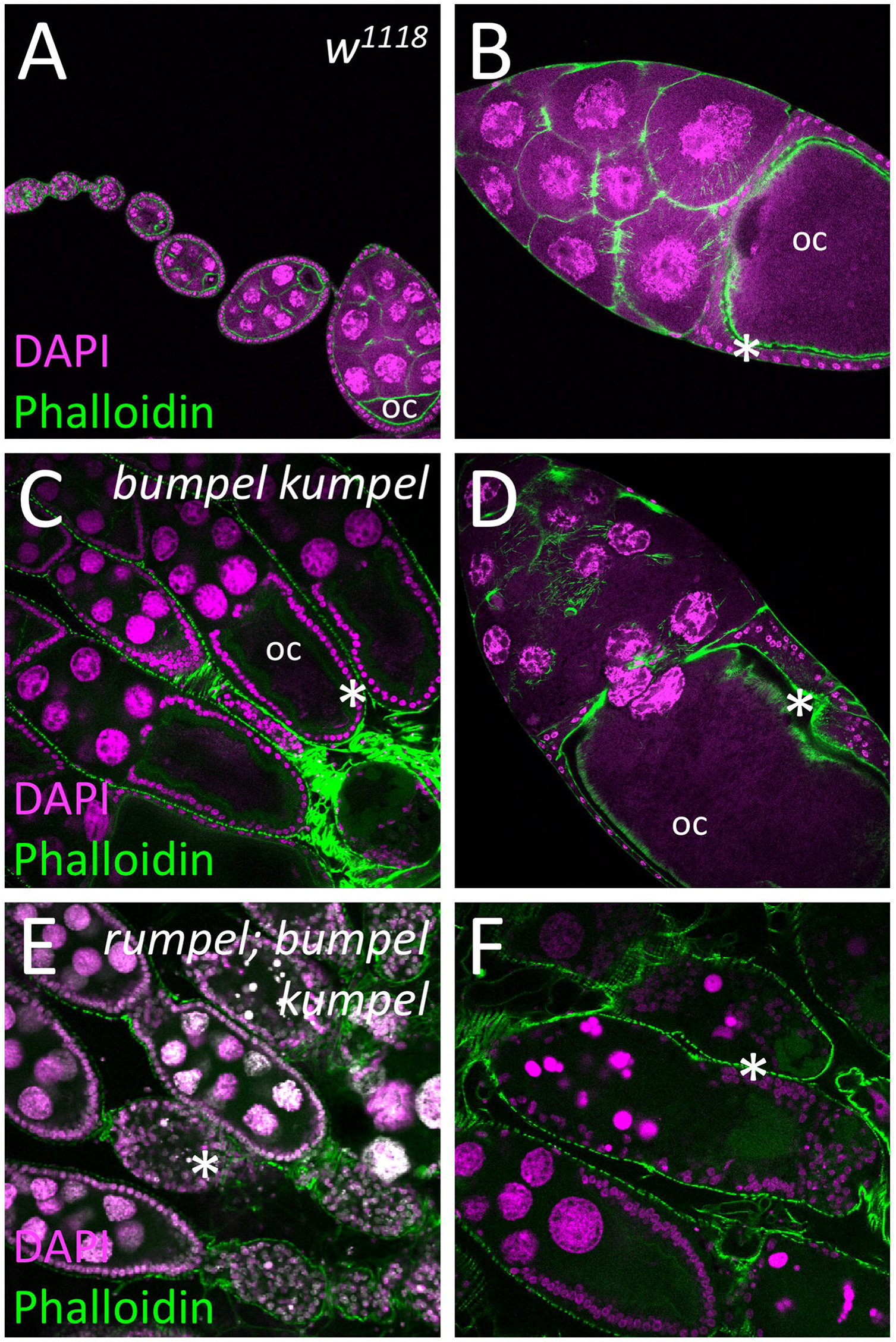
SLC5A transporters are required for the oogenesis. Confocal analysis of wild type and mutant ovaries. Nucleii are laballed by DAPI staining, F-actin is shown following phalloidin staining (green). **(A,B)** In control females oogenesis developing egg chambers connected by stalk cells mature to form tubular ovarioles. During the previtellogenic phase, the future oocyte (oc) is defined which is positioned at the posterior pole. During the vitellogenic phases the oocyte grows exponentially and is surrounded by a cuboidal follicular epithelium (asterisks). **(C,D)** Homozygous *bumpel kumpel* double mutants are sterile but lay few eggs. Oogenesis is affected at the vitellogenic phase. The oocyte and the follicle epithelium degenerate. **(E,F)** Homozygous *rumpel bumpel kumpel* mutants are sterile and never lay eggs. Oogenesis is affected at the vitellogenic stage as seen in *rumpel bumpel* double mutants. However, the disintegration of oocytes and the follicular epithelium is more pronounced.

We next tested whether the triple mutant shows a locomotor phenotype. At 25°C, *rumpel^Δ+cherry^* mutant larvae have slightly reduced distance to origin after 3 minutes free crawling compared to control larvae (**supplementary Fig. 4**). A very similar reduction in the distance to origin is detected for the triple mutant (**supplementary Fig. 4**). In contrast, at elevated temperature (32°C) the triple mutant shows a significantly reduced distance to origin whereas control and *rumpel^Δ+cherry^* mutants are not affected by the increase in temperature (**supplementary Fig. 4**). Further analysis of the different locomotion parameters revealed that although triple mutant larvae have a reduced distance to origin, they are faster, but show an altered bending behavior (**supplementary Fig. 4**).

### *rumpel, bumpel* and *kumpel* encode SLC5A homologs that likely transport lactate

The sterility phenotype shown by the double mutant animals allowed us to conduct rescue experiments. One copy of the *bumpel* minigene rescued fertility of the *bumpel kumpel* double mutant. Interestingly, overexpression of *bumpel* by introducing two copies of the *bumpel* minigene into a wild type background but not in a heterozygous mutant background causes a lethal phenotype. Likewise, pan-glial *repo-Gal4* based overexpression of a *UAS-Bumpel^3xHA^* construct (Bischof et al., 2013) causes lethality. Rescue experiments using Gal4 based expression of the *kumpel^PD^* isoform resulted in a few larvae, but did not rescue beyond larval stages. This suggests that expression levels are likely crucial for function.

The predicted Rumpel, Bumpel and Kumpel proteins all belong to the SLC5A family of monocarboxylate transporters that utilize a sodium gradient across the plasma membrane to transport a variety of solutes. To determine the nature of these solutes we utilized the knowledge on transported metabolites of the mammalian orthologues SLC5A1-SLC5A12. We obtained full-length cDNAs, encoding all transporters except SLC5A4 and SLC5A6 (**Table 1**) and generated transgenic flies expressing the different mammalian SLC5A proteins under UAS control. We generated females with the following genotypes [*Act5C-Gal4/UAS-SLC5Axy*, *bumpel kumpel* / *bumpel kumpel*] and assayed whether female sterility was rescued when crossed to CantonS males. To our surprise, almost all transgenes showed very limited rescue after prolonged culture (**Table 1**). Ubiquitous expression of SLC5A2, which transports glucose (Wright, 2013), gave robust rescue with many flies eclosing from homozygous *bumpel kumpel* mothers. Weaker rescue with low numbers of eclosing flies was noted following expression of SLC5A8 and SLC5A12 (which transport lactate (Coady et al., 2004; Gopal et al., 2007; Miyauchi et al., 2004)) (**Table 1**).

**Table 1.** Mammalian SLC5A proteins can compensate the function of Drosophila orthologs. The ability of mammalian different SLC5A genes indicated to rescue the sterility phenotype associated with the *bumpel kumpel* double mutant is given. The predicted solutes transported by the different carriers is indicated (Wright, 2013). The strength of the rescuing ability is indicated by +. Flies indicate full rescue and flies appeared. The Gal4 driver used in the respective rescue experiments is indicated.

| Gene | Protein | Predicted Solute | Species | Rescue | Gal4 driver |
| --- | --- | --- | --- | --- | --- |
| *SLC5A2* | SGLT2 | Glucose | *M. musculus* | +++ flies | *act5C-Gal4* |
| *SLC5A8* | SMCT1 | Lactate, pyruvate | *M. musculus* | ++ flies | *act5C-Gal4* |
| *SLC5A12* | SMCT2 | Lactate, nicotinate | *H. sapiens* | ++ flies | *act5C-Gal4* |
| *SLC5A1* | SGLT1 | Glucose, galactose | *M. musculus* | + flies | *act5C-Gal4* |
| *SLC5A7* | CHT | Choline | *M. musculus* | + flies | *act5C-Gal4* |
| *SLC5A11* | SMIT2 | Glucose, myo-Inositol | *R. norvegicus* | + flies | *act5C-Gal4* |
| *SLC5A3* | SMIT1 | myo-Inositol | *M. musculus* | + | *act5C-Gal4* |
| *SLC5A5* | NIS | Iodite | *M. musculus* | + | *act5C-Gal4* |
| *SLC5A9* | SGLT4 | Mannose, fructose | *M. musculus* | + | *act5C-Gal4* |
| *SLC5A10* | SGLT5 | Mannose, fructose | *M. musculus* | + | *act5C-Gal4* |
| *SLC5A2* | SGLT2 | Glucose | *M. musculus* | + | *repo4.3-Gal4* |
| *SLC5A2* | SGLT2 | Glucose | *M. musculus* | + flies | *nrv2-Gal4* |
| *kumpel* | Kumpel<sup>PD</sup> |  | *D. melanogaster* | + | *act5C-Gal4* |
| *SLC5A4* | SGLT3 | Na<sup>+</sup> / glucose sensor | *M. musculus* | nd |  |
| *SLC5A6* | SMVT | Multi-vitamine, biotin, lipotate | *M. musculus* | nd |  |

The above rescue data suggests that ubiquitously induced glucose and/or lactate transport is able to rescue the *kumpel bumpel* double mutant sterility phenotype. To address the question whether glial expression is sufficient for rescue, we expressed SLC5A2, SLC5A8 and SLC5A12 using *repo-Gal4* and *nrv2-Gal4*. Interestingly, panglial expression resulted in only a few, small larvae without any eclosed flies; whereas rescue using *nrv2-Gal4* directed expression produced a small number of surviving flies.

## Discussion

Here we describe the analysis of three predicted Drosophila solute carrier proteins: Rumpel, Kumpel and Bumpel; which show overlapping expression patterns in the larval and the adult CNS. The *rumpel* gene was initially identified in an RNAi-based screen for adult locomotor defects (Schmidt et al., 2012). The same behavioral phenotype was also found in a different study but unlike our findings the phenotype was assigned to defects in astrocyte-like glial cells (Ng and Jackson, 2015). We thus generated mutants and surprisingly, detected only very weak behavioral phenotypes. The notion that the locomotor phenotype was stronger in animals carrying a CRISPR induced point mutant compared to animals carrying a deletion of the *rumpel* locus could possibly indicate the presence of a nonsense mediated decay mechanism that leads to a more global change of the transcriptional activity of the cell. This was also corroborated by the finding that flies lacking all ensheathing glia show a pronounced heat shock sensitivity.

To determine whether a possible genetic redundancy is causing this apparent lack of abnormal phenotypes in *rumpel* mutants, we analyzed the most closely related genes *kumpel* and *bumpel*. However, even triple mutants do not show the initially observed RNAi-induced locomotor phenotype; though we did find a prominent reduction in exploratory locomotion and thus a reduced distance to origin. A genetic redundancy, however, is detected during oogenesis. In contrast to single mutants, in *bumpel kumpel* double mutants, oogenesis is defective during the vitellogenic phase and in *rumpel bumpel kumpel* triple mutant females oogenesis arrests shortly before the vitellogenic phase.

In Drosophila, the complexity of the SLC5A family transporters is similar to the one found in mammals and 14 SLC5A proteins are encoded in the fly genome (Featherstone, 2011). Only three of these genes have been analyzed in greater detail. The *Sodium-dependent multivitamin transporter* (*Smvt*) is not expressed in the central nervous system but its muscle specific knockdown causes a flightless phenotype (Schnorrer et al., 2010). The gene *salty dog* (*salt*) is also not expressed in the CNS and affects survival in a high-salt environment (Stergiopoulos et al., 2009). The gene *cupcake* (*SLC5A11*) is prominently expressed in neurons of the adult fly brain where it is required for proper food selection (Dus et al., 2013). Here, we present analysis of three additional members of the SLC5A family which not only share protein homology but also show overlapping expression patterns. All single mutants are viable and fertile with no obvious behavioral phenotypes.

Similar to *rumpel*, where a RNAi mediated knockdown causes a phenotype but not the *rumpel* mutant, *Mfs3* and *pippin* knockdown but not *Mfs3* and *pippin* mutants show a compensatory upregulation of Tret1-1. Interestingly, *Mfs3*, *pippin* and *Tret1-1* all encode carbohydrate transporters. Which mechanisms could account for such effects? Most easily they could be explained as off-target effects, however, no such off-targets are predicted for the RNAi strain used (Dietzl et al., 2007). An alternative explanation is that the construct directing expression of double RNA is inserted in gene that dominantly contributes to the phenotype. However, it is also possible that this is a result of a more general transcriptional adaptation (Sztal and Stainier, 2020). First identified in zebrafish while studying the role of the endothelial extracellular matrix (ECM) protein *Egfl7* (Rossi et al., 2015). Morpholino-based knockdown of the gene encoding this protein resulted in a severe phenotype in zebrafish (as well as in Xenopus or in human cells) but the corresponding mutant appeared largely normal. Subsequent mass spectrometry revealed the upregulation of the related ECM protein Emilin3a in the *Egfl7* mutant but not in the knockdown animal. Subsequently it was shown that mutant mRNA degradation plays a crucial role in activating transcriptional adaptation in zebrafish and mouse cell lines and a clean deletion of the gene is not sufficient to trigger transcriptional adaptation (El-Brolosy et al., 2019). However, in contrast to what is observed for zebrafish, *rumpel* excision mutants that lack all *rumpel* transcripts still exhibit no abnormal phenotype.

Here we show that Rumpel, Kumpel and Bumpel are all expressed in overlapping sets of CNS glial cells. In addition, RNA-seq data obtained from dissected tissues show no expression of *rumpel*, *bumpel* or *kumpel* in ovaries (Graveley et al., 2011). Yet, the double and triple mutants are sterile with an ovary phenotype, which allowed us to perform rescue experiments using an array of heterologous SLC5A transporters with defined solute transport properties. As control, we performed rescue using the *bumpel^minigene^* construct. Homozygous *bumpel^minigene^* animals are lethal and this lethality is rescued by the *bumpel kumpel* double mutant. Likewise, the sterility phenotype associated with homozygous *bumpel kumpel* double mutants is rescued by a single copy of the *bumpel^minigene^*. Thus, the expression level of *bumpel* appears tightly regulated, and an overexpression of SLC5A is more detrimental than a complete lack of SLC5A transporter. This may have obscured the rescue experiments using heterologous SLC5A sequences, which indicated that Bumpel and/or Kumpel transport glucose or lactate.

The Drosophila adult ovary comprises a pair of 16-20 tubular ovarioles, where egg chambers connected by stalk cells mature in a sequential manner (Margolis and Spradling, 1995; Robinson and Cooley, 1997; Spradling, 1993). Two phases of egg chamber growth can be defined: the previtellogenic and the vitellogenic phases. During the vitellogenic phase an exponential growth of the oocyte occurs due to yolk import through the follicular cells surrounding the egg chamber. The onset of the vitellogenic phase is controlled by hormones and the nutritional state of the fly which is generally regulated by insulin-like peptides (Mendes and Mirth, 2016; Mirth et al., 2014; Nässel and Vanden Broeck, 2016; Raushenbach et al., 2004; Richard et al., 2005). The *rumpel bumpel kumpel* triple mutant specifically affects the vitellogenic phase, suggesting that glucose metabolism in the brain controls ovary development. Such a brain-gonad axis had been described before. For example the neurotransmitter octopamine, which is closely related to norepinephrine, is known to act as an alerting signal in insects and octopaminergic neurons reach almost all peripheral tissues (Pauls et al., 2018). The endocytic regulator monensin-sensitive 1 (Mon1) is required in octopaminergic neurons for normal ovary growth and a cell-type specific knockdown results in the absence of late stage egg chambers (Dhiman et al., 2019). Octopamine also reaches the insulin producing cells (IPCs) which are known to regulate feeding behavior and express the octopamine receptor OAMB1 (Luo et al., 2014; Selcho and Pauls, 2019). It is possible, disruption of SLC5A function in the Drosophila glial cells affects nutrient sensing in the nervous system, which feeds back to octopaminergic neurons, and thereby hinders ovarian development.

## Materials and Methods

### Drosophila work

Unless otherwise stated, all Drosophila stocks and crosses were raised on standard Drosophila food at 25°C. To target all glial cells we employed *repo4.3-Gal4* (Schmidt et al., 2012), to specifically target glial subsets we used *83E12-Gal4* (ensheathing glia) (Li et al., 2014; Otto et al., 2018), *nrv2-Gal4* (cortex and ensheathing glia) (Sun et al., 1999), R55B12-Gal4 (cortex glia) (Li et al., 2014), *alrm-Gal4* (astrocyte-like glia) (Muthukumar et al., 2014). UAS-dsRNA lines targeting *rumpel* (GD3270, KK106220), *Orco* (KK100825) were obtained from the VDRC (Vienna, Austria). *UAS-GFP^dsRNA^* (BL9330, BL9331), *kumpel^MiMIC-GFSTF^* (BL60231), *UAS-lam∷GFP* (BL7378), *Act5C-Gal4* (BL4414), *UAS-rpr.C* (BL5823), *UAS-hid.Z* (BL65403), and *Canton S* (BL64309) were obtained from the Bloomington stock center (Bloomington, Indiana, USA). *UAS-bumpel* (F003123) was obtained from the FlyORF collection (Zürich, Switzerland).

### Generation of mutants and transgenes

All single-guide RNA (sgRNA) and PCR primers used in this study are listed below in Table 2. To generate mutants, sgRNA plasmids were injected into Cas9 expressing recipient embryos (Port et al., 2014). Indels were detected in the F1 generation using PCR and subsequent sequence analysis. To replace the *rumpel* locus with mCherry, we generated a donor plasmid starting from pTV3 (kindly provided by J.P. Vincent, London) where mCherry is flanked by 2 kb flanking genomic *rumpel* DNA. To generate endogenous V5-tags, we generated donor plasmids where the V5 tag is inserted just before the stop codon, flanked by 1.5 kb of genomic sequence on either side. The *bumpel^V5^* minigene spans ~3.1 kb of genomic DNA, including the UTRs of the neighboring genes. We inserted V5 tag just before the stop codon using standard procedures. The construct was inserted into the *44F* landing site (Bischof et al., 2007). The different mutants and transgenes generated in this study are listed in Table 3.

**Table 2.**
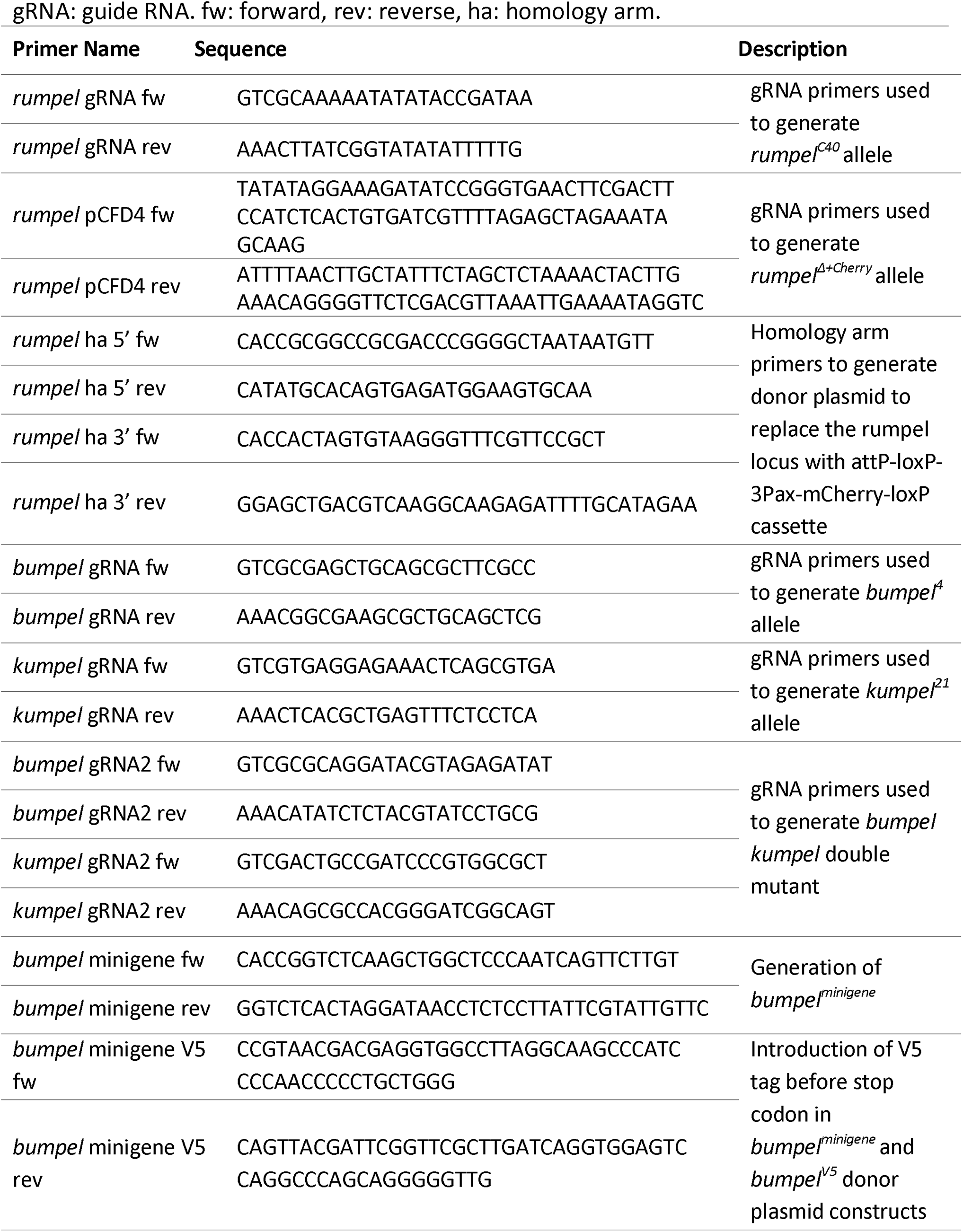
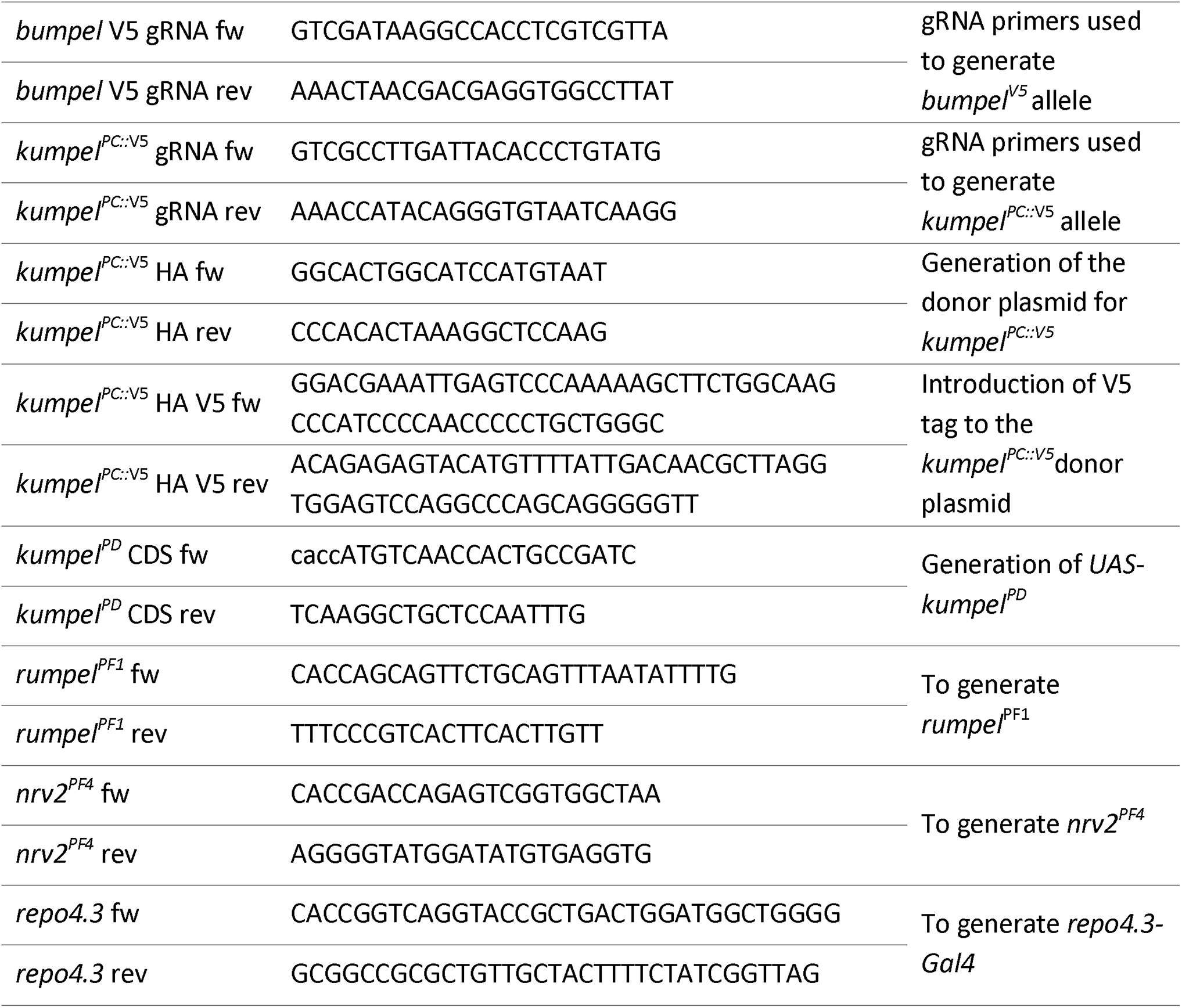
Primers used in this study.

**Table 3.**
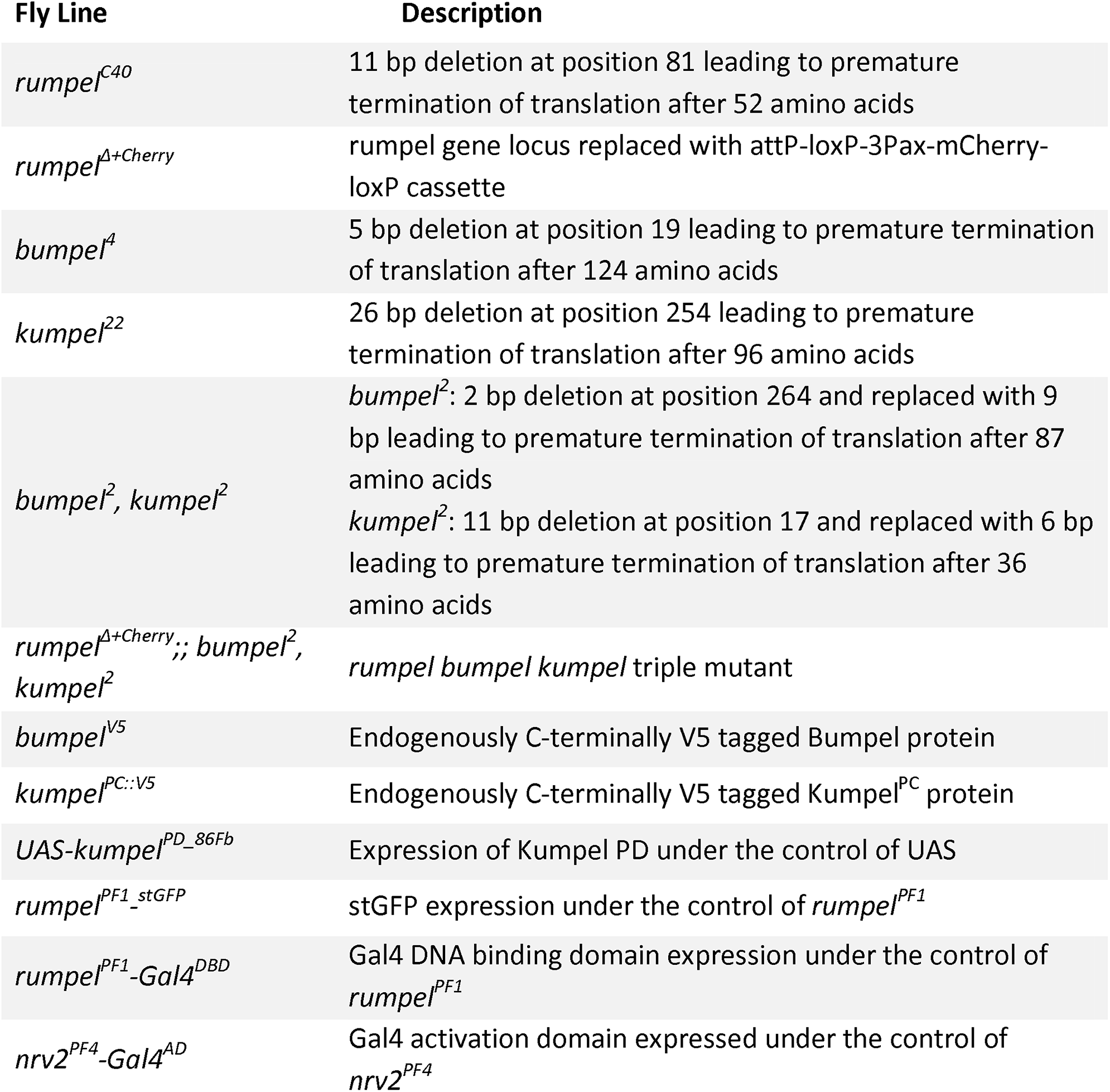
Mutant and insertions generated in this study.

### Generation of Interspecies rescue construct

To generate interspecies rescue constructs, we amplified the open reading frames of the mammalian *SLC5A* genes from their respective cDNA plasmids which were obtained from Thermo Fisher (see Table 4). We generated UAS-based expression plasmids using *pUAST-attB-rfa* and inserted them into *51C^RFP+^* landing site.

**Table 4.** cDNA clones used to generate transgenic fly strains expressing mammalian SLC5A genes.

| Name | cDNA clone | Accession No. | Species |
| --- | --- | --- | --- |
| *Slc5a1* | MMM1013-202761995 | BC003845 | *Mus musculus* |
| *Slc5a2* | MMM1013-202765784 | BC022226 | *Mus musculus* |
| *Slc5a3* | EMM1002-213343621 | BC140982 | *Mus musculus* |
| *Slc5a5* | EMM1002-213341390 | BC137650 | *Mus musculus* |
| *Slc5a7* | MMM1013-202859819 | BC065089 | *Mus musculus* |
| *Slc5a8* | MMM1013-202765927 | BC017691 | *Mus musculus* |
| *Slc5a9* | MMM1013-202767795 | BC021357 | *Mus musculus* |
| *Slc5a10* | MRN1768-202721502 | BC161867 | *Rattus norvegicus* |
| *Slc5a11* | MMM1013-202765603 | BC031742 | *Mus musculus* |
| *SLC5A12* | MHS1010-202801048 | BC049207 | *Homo sapiens* |

For the interspecies rescue experiments, we have recombined *Act5C-Gal4* with *UAS-SLC5Axy* and then generated *Act5C-Gal4*, *UAS-SLC5Axy*; *bumpel^2^*, *kumpel^2^*. Generated homozygous females were crossed with wild type CantonS males. The fertility rescue was scored as larval, pupal, full rescue or no rescue in the F1 generation. Furthermore, *UAS-SLC5A2* was recombined with *repo4.3-Gal4* and *nrv2-Gal4*. Glial interspecies rescue was performed as explained above.

### Behavioral analyses

Larval behavioral experiments were performed at 25°C unless otherwise indicated. Larval locomotion was analyzed using FIM (Risse et al., 2013; Risse et al., 2017a). Locomotion of 10-15 larvae was recorded for 3 min at 10 frames per second. Tracking data was analyzed as described (Otto et al., 2018; Otto et al., 2016; Risse et al., 2014). Distance to origin is defined by the distance of the larva from the spot where it placed on the agar plate, normalized per minute. In bending distribution plots, the number of head bends per 10 seconds deviating from the larval 180° body axis are shown. Velocity during Go-phases is defined as distance per time (pixels per second) during larval go phases.

To obtain a heatmap representation of larval distribution on the tracking area we employed the open source RStudio software (http://www.rstudio.com/). In a custom-made script, the tracking area was divided into 50×50 px squares which is in the same size range as the average larval length (45 px in these settings). The frequency of an appearance of a larva in each square was calculated and is indicated by shading intensity. In all experiments, the same number of larvae in the same area over the same length. For the analysis of the sleep phenotype, freshly hatched males were collected and aged at 25°C for 3 days. 40 single 3-days old male flies placed into individual capillaries mounted in an Ethoscope arena were analyzed as described (Geissmann et al., 2017). Here individual flies are constantly video-tracked for 7 days at constant temperature (25°C) and humidity (65%) with 12h light-dark cycle. Sleep was defined as 5 minutes with no activity and was quantified using the Rhetomics package in R (Geissmann et al., 2019). For statistical analyses the standard non-parametrical Wilcoxon rank-sum test was performed using FIM Analytics 2 or R.

To test for temperature sensitivity, 3 - 5 days old flies staged males and females were collected and 5 flies each were placed into a new vial. On the next day, the 5 flies were transferred into a fresh empty vial without food and anesthesia. After a 5-10 min acclimation time the vials were placed in a water bath at 40°C for 2 min. Afterwards, the vials were filmed for 4 min at room temperature. Flies, which lay on their backs and did not move, were counted as paralyzed. 100 flies for each genotype were recorded.

To determine the negative geotaxis, 100 females were separately collected and staged as required. 10 flies each were placed in fresh vials with standard food and kept overnight at 25 °C. Afterwards, the 10 flies were loaded in long plastic tubes without anesthesia. After 5-10 min acclimation time the tubes were placed in the rapid iterative negative geotaxis (RING) system according to Gargano et al., 2005. In total, 100 flies were tested. The images were processed using Fiji with MTrack3_.jar plugin and AutoRun2.ijm macro. The mean velocity was determined using the RING assay Script.R in the R program. Statistical analysis was performed by Prism 6.0.

### Immunohistochemistry

Immunohistochemistry was performed according to standard protocols except the fixation process. Larvae were dissected in ice-cold PBS. Filet preparations were fixed in Bouin’s Solution (Sigma-Aldrich) for 3 min at room temperature, while the adult brains were fixed in 2% PFA for 90 min at room temperature. The following antibodies were used: mouse anti-Repo (DSHB, 1:5), rat anti-DN-Cadherin (DSHB, 1:10), mouse anti-V5 (Thermo Fisher, 1:500), rabbit anti-GFP (Thermo Fisher, 1:1000), rabbit anti-dsRed (Clontech, 1:1000), goat anti-HRP-Cy5 (Dianova, 1:200). Rabbit anti-Rumpel peptide antibodies were generated against the C-terminal domain of Rumpel (Pineda, Berlin, 1:500). Secondary antibodies conjugated to Alexa Fluor 488, Alexa Fluor 568 or Alexa Fluor 647 were used (Thermo Fisher, 1:1000,). Ovaries were dissected in ice-cold PBS and fixed in 4% PFA for 15 min. Immunohistochemistry for ovaries was performed as described (Bogdan et al., 2005). Alexa Fluor 568 Phalloidin (Thermo Fisher, 1:100) and DAPI (Thermo Fisher, 1:1,000). Confocal microscopy data was generated using a Zeiss 710 or 880 LSM or Leica TCS SP8 DLS. Images were acquired using either the Zeiss LSM ZEN imaging software, or the LSM LAS X software and analyzed using Fiji (Schindelin et al., 2012).

### Paralogs and orthologs of *rumpel*

The amino acid sequences of Rumpel and its closest mouse orthologues SLC5A5, SLC5A8 and SLC5A9 were aligned and visualized using T-Coffee tool (http://tcoffee.crg.cat/apps/tcoffee/do:regular) and Boxshade (https://embnet.vital-it.ch/software/BOX_form.html). For amino acid sequence comparisons, we used Clustal Ω (https://www.ebi.ac.uk/Tools/msa/clustalo/) and Blast (https://blast.ncbi.nlm.nih.gov). To reconstruct the phylogenetic tree of the *rumpel* paralogs we employed MEGA X (default settings, (Kumar et al., 2018; Stecher et al., 2020)).

## Acknowledgements

We are thankful to M. Ogueta for help during the sleep analysis, B. Altenhein and J.P. Vincent for sharing antibodies and plasmids. We are thankful to E. McMullen, J. Bittern for help in analysis of larval locomotion and all members of the Klämbt lab for help throughout the project.

## Author contributions

Conceptualization: K.Y., C.K.; Methodology and investigation: K.Y., S.S., S.T., N.P., B.W., M.B.; Software: L.G.; Writing - original draft: K.Y., C.K.; Writing - review & editing: K.Y., C.K., B.W. M.B.; Visualization: K.Y., L.G., M.B., N.P., C.K.; Funding acquisition: C.K.

## Data availability

All reagents are available from the corresponding author upon reasonable request.

